# Rapid and cost-effective epitope mapping using PURE ribosome display coupled with next-generation sequencing and bioinformatics

**DOI:** 10.1101/2023.09.09.556969

**Authors:** Beixi Jia, Teruyo Ojima-Kato, Takaaki Kojima, Hideo Nakano

## Abstract

A novel, efficient and cost-effective approach for epitope identification of an antibody has been developed using a ribosome display platform. This platform, known as PURE ribosome display, utilizes an *Escherichia coli*-based reconstituted cell-free protein synthesis system (PURE system). It stabilizes the mRNA-ribosome-peptide complex via a ribosome-arrest peptide sequence. This system was complemented by next-generation sequencing (NGS) and an algorithm for analyzing binding epitopes. To showcase the effectiveness of this method, selection conditions were refined using the anti-PA tag monoclonal antibody with the PA tag peptide as a model. Subsequently, a random peptide library was constructed using 10 NNK triplet oligonucleotides via the PURE ribosome display. The resulting random peptide library-ribosome-mRNA complex was selected using a commercially available anti-HA (YPYDVPDYA) tag monoclonal antibody, followed by NGS and bioinformatic analysis. Our approach successfully identified the “DVPDY” sequence as an epitope within the hemagglutinin amino acid sequence, which was then experimentally validated. This platform provided a valuable tool for investigating continuous epitopes in antibodies.

## Introduction

Antibodies are a class of proteins capable of identifying and binding to specific molecules, with broad implications in scientific research, diagnostics, and therapeutic applications [1]. Monoclonal antibodies, which bind selectively to target molecules, are integral in addressing a variety of diseases, encompassing autoimmune conditions, infectious pathogens, and cancer [2]. To accommodate the growing demand for monoclonal antibodies, a range of screening methods have been devised. One of these methods is Ecobody Technology [3], which is a rapid and cost-effective method for screening and producing monoclonal antibodies from individual animal B cells.

An epitope, a discrete region of an antigen recognized and bound by an antibody [4], dictates the function of each respective antibody. Therefore, epitope identification or ‘epitope mapping’ is a crucial step in characterizing the procured monoclonal antibodies [5]. While several methodologies exist for epitope mapping, all experimental techniques are dependent upon the physical interaction between the antibody and the antigen [6]. X-ray crystallography [7], an X-ray analysis method applied to an antigen-antibody crystal complex, can provide high-resolution structural information of the epitope. It is widely regarded as the most informative and trustworthy method for epitope mapping. Another approach to obtain the conformational information about the epitope is the deuterium exchange method combined with Mass Spectrometry (MS) analysis [8]. However, both approaches require substantial amounts of pure antibody and antigen, which necessitate time-consuming and costly experiments.

An alternative approach for epitope mapping involves the use of a peptide microarray [9]. This microarray consists of overlapping peptides that span the full length of the target protein and are immobilized on a solid substrate. The antibody of interest is bound to this array and subsequently detected via fluorescence or chemiluminescence to define its epitope. However, creating a custom peptide array for each antigen is relatively expensive, and a unique array is needed for each different antigen.

Alternatively, a peptide library generated by phage display [10] or *in vitro* display technologies can also be employed for epitope mapping. The *in vitro* display technologies establish a physical linkage between the phenotype (protein or peptide) and the genotype (mRNA or cDNA) via either a ribosome (in ribosome display [11]) or the puromycin (in mRNA or cDNA display [12–13]). In 2005, Schimmele and co-workers successfully implemented epitope mapping against the Nogo Receptor using ribosome display with an *E. coli* S30 extract [14]. In 2009, Osada et al. undertook epitope mapping of an anti-β-cat monoclonal antibody utilizing ribosome display with an *E. coli*-based reconstituted cell-free protein synthesis system (named as PURE-ribosome display system) [15]. This system, having negligible amounts of protease and nuclease [16], enhances the stability of the peptide-ribosome-mRNA complex. Traditional Sanger sequencing has been used to analyze the results, but this method is laborious and limits the number of sequences that can be studied. More recently, Next-Generation Sequencing (NGS) has been integrated with in vitro display technologies to monitor sequence selection more efficiently [17–18].

In this study, after optimization of the PURE-ribosome display condition utilizing anti-PA tag antibody, we developed a rapid and cost-effective epitope mapping platform using ribosome display using the commercially available PURE cell-free protein synthesis system, followed by the NGS and an in-house Python program (Figure 1A).

**Figure 1.**
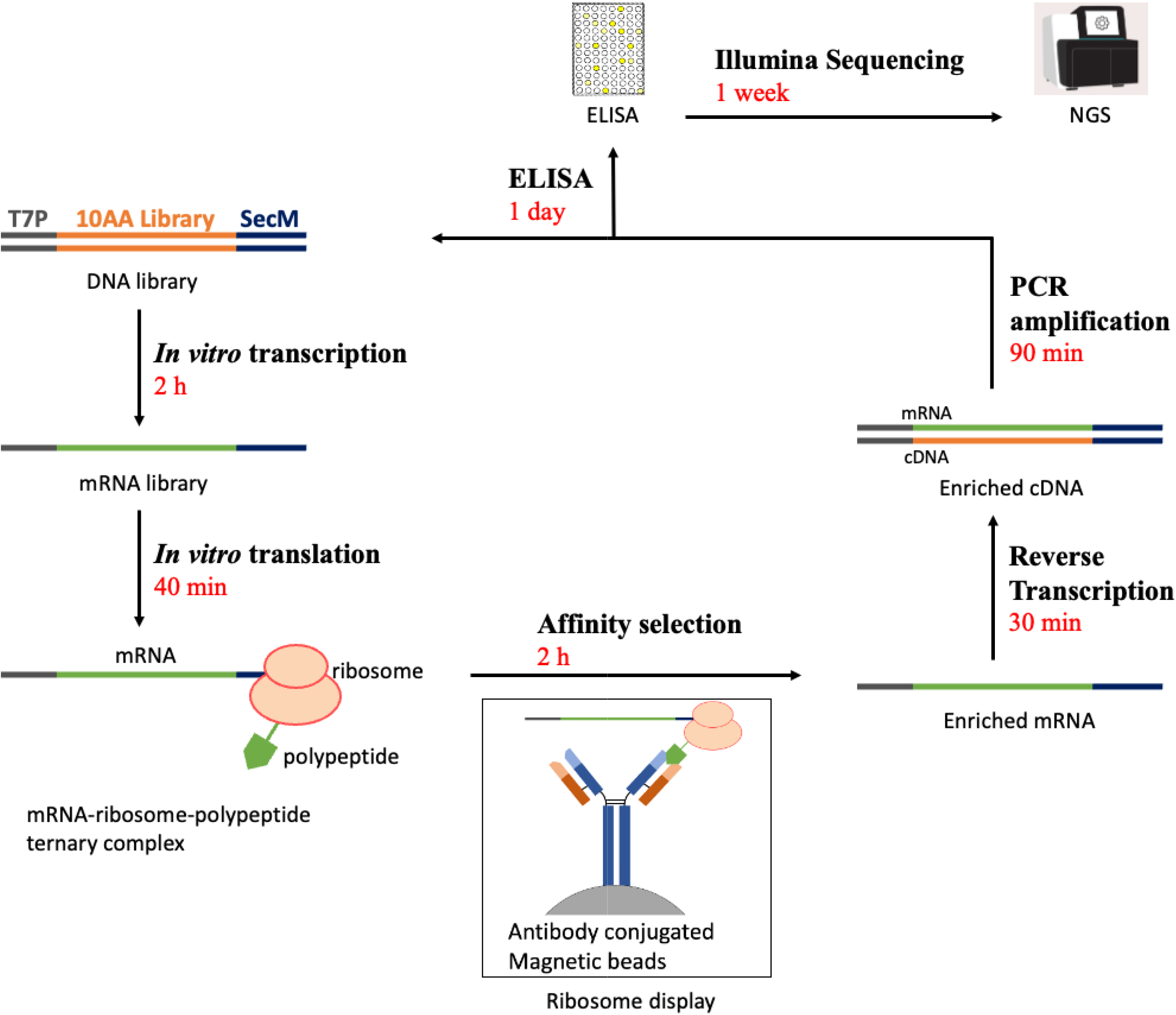
The overview of PURE ribosome display platform. (A) Schematic protocol of the PURE ribosome display platform. The library was constructed and subjected to *in vitro* transcription. The mRNA was translated using the PURE*frex* 1.0 kit, and the resulting mRNA-ribosome-polypeptide ternary complex underwent affinity selection with antibody-conjugated magnetic beads. Following disassociation with EDTA, the enriched mRNA was reverse-transcribed and PCR-amplified to prepare the template for the next round of selection. Concurrently, the PCR-amplified products were transcribed and translated for ELISA to assess binding affinity against the target antibody, and the products with an increasing ELISA signal were sequenced using the Illumina Nextseq 550 system. The preparation times for each step were indicated in red. (B) The construction of the DNA Library for ribosome display. The DNA library comprised a T7 promoter, the Ribosome Binding Site, the T7 tag, SK sequence, 10-codon “NNK” triple oligonucleotide random library, the portion of protein D, TEV site, and portion of SecM arrest sequence. GS linker indicated the amino acid sequence of “GGGS”.

## Materials and Methods

### PURE Ribosome Display for Model Binary Library against anti-PA Tag Antibody

To establish model binary library for ribosome display selection against anti-PA tag antibody, plasmids containing His tag, either of PA tag (GVAMPGAEDDVV) [19] or Flag tag (DYKDDDDK), protein D (T20-V109) [20–21], and SecM arrest sequence (A148-T170) [22–23] were constructed. The oligo DNA primers used in this experiment were procured from Eurofins, Japan (refer to Table S1 for details). These constructed plasmids served as templates for PCR amplification using the F1 primer and the Fag-Rev primer. The amplified products were then purified using a spin column (Nippon Genetics Co.Ltd, Japan) and were used as a control-binder (PA tag) or a non-binder (Flag tag) for ribosome display selection.

The purified DNAs were used as templates for *in vitro* transcription, conducted with the RiboMAX Large Scale RNA Production System-T7 (Promega, USA) at 37℃ for 120 minutes. This was followed by a 30-minute DNase I (Takara Bio Inc., Japan) digestion at 37℃. The synthesized mRNA was subsequently purified using a NucleoSpin® RNA Clean-Up Kit (Takara Bio Inc., Japan). The concentration of the purified mRNA was quantified using a Nanodrop One (Thermo Scientific, USA).

The resulting mRNA was mixed in a control binder: non-binder at ratio of either 1:10 (5 ng:50 ng) or 1:100 (1 ng:100 ng). The mixed mRNAs were then incorporated into a 20 µL-scale reaction of the cell-free protein synthesis (CFPS) system PURE*frex*1.0 (GeneFrontier, Japan) and incubated at 37℃ for 25 minutes. The reaction mixture was subsequently placed on ice for 2 minutes, followed by the addition of 100 µL of ice-cold stop buffer (50 mM Tris-HCl, pH 7.5; 150 mM NaCl; 50 mM Mg(OAc)_2_; 0.1%(v/v) Tween-20; 2.5 mg/mL sodium heparin) to halt the reaction [16]. The mixture was centrifuged at 4℃ for 10 minutes to remove insoluble proteins, and the supernatant was prepared for ribosome display selection.

The supernatant was incubated with 50 µL of anti-PA tag antibody beads (FUJIFLIM Wako Pure Chemical Corporation, Japan) at room temperature for 1 hour. The beads were then washed four times with a washing buffer (50 mM Tris-HCl, pH 7.5; 150 mM NaCl; 50 mM Mg(OAc)_2_). The mRNA bound to the beads was dissociated by adding an elution buffer (50 mM Tris-HCL, pH 7.5; 150 mM NaCl; 50 mM EDTA; 10 µg/mL total RNA from yeast (Sigma, USA)) [24] and rotating at room temperature for 20 minutes. The collected supernatant was then subjected to reverse transcription using SuperScript Ⅳ reverse transcriptase (Thermo Fisher, USA) and the RT-Rv primer. Since the elution buffer contains EDTA, which has the potential to inhibit the enzymatic activity, Mg(OAc)_2_ buffer was added at a final concentration of 50 mM during the reverse transcription step. The transcribed cDNA was amplified using a nested two-step PCR amplification process employing KOD One PCR Master Mixture (TOYOBO, Japan) along with the PCR1-Fw/Rv and the PCR2-Fw/Rv primers. The amplification was conducted under the following thermal conditions: initial denaturation at 94℃ for 3 minutes, followed by 25 cycles of denaturation at 98℃ for 10 seconds, annealing for 5 seconds at 56℃ for the first PCR step or 61℃ for the second PCR step, and extension at 68℃ for 10 seconds.

To differentiate between the non-binder (Flag tag) and the control-binder (PA tag) DNA, a unique BamHⅠ restriction site, present only in the non-binder DNA, and absent in the control-binder DNA, was utilized (Figure S1). The PCR products from the control-binder, the non-binder, the input binary libraries (1:10 or 1:100) and the selected pools were subjected to restriction digestion using BamHI (Takara Bio Inc., Japan). The digestion products were then analyzed using a 10% polyacrylamide gel electrophoresis (PAGE) running at 10 mA, 100 V for a duration of 70 minutes.

### Preparation of random nucleotide library for PURE ribosome display

Initially, a backbone plasmid named RD backbone was constructed using HiFi DNA Assembly. This plasmid included an upstream T7 promoter for *in vitro* transcription, a ribosome binding site (RBS) for *in vitro* translation, a segment of the SKIK sequence for high-level expression [25], a T7 tag for detection, a spacer sequence from bacteriophage Lambda head protein D to further enhance expression [20–21], and the SecM arrest sequence, which stalled the ribosome and formed a ternary complex [22–23]. It was subsequently amplified through PCR to create the library vector with theLibvec-For and Libvec-Rev primers, followed by purification with spin column.

A random 10 NNK DNA library, which represented all 20 distinct amino acids, was synthesized by mixing an oligo DNA (10AA-Fw) containing 10 NNK triplet oligonucleotides and the Lib-Rev primer with dNTPs and Klenow buffer (Takara Bio Inc., Japan) for heating at 95℃ for 3 minutes and cooling at room temperature for 5 minutes. Klenow Fragment (Takara Bio Inc, Japan) was then added to the mixture, followed by incubating at 1 hour at 37℃, and a 5-minute inactivation at 65℃. The library DNA fragment was then ligated to the library vector utilizing NEBuilder HiFi DNA Assembly (New England Biolabs, Japan) at 50℃ for 15 minutes. The assembled products were used as template for PCR amplification to obtain the input random library, as shown in Figure 1B, for ribosome display using F1 and Fag-Rev primer,

### PURE ribosome display selection of random library

The target antibody, an anti-HA tag monoclonal antibody (MBL Co. Ltd., Japan) was biotinylated using the Biotin Labeling Kit-NH_2_ (Dojindo Laboratories Co., Ltd., Japan), and subsequently bound to Dynabeads M-280 Streptavidin (Invitrogen Thermo Fisher Scientific, USA) at room temperature for 30 minutes.

The cell-free protein synthesis (CFPS) solution supernatant was prepared as described similarly to that in the binary model library using PURE*frex*1.0 and 150 ng mRNA synthesized from the input random library DNA as template. To minimize non-specific binding, the CFPS solution was first incubated with magnetic beads without ligand for 30 minutes at room temperature. The residual supernatant was then incubated with antibody-immobilized magnetic beads for 1 hour at room temperature. The subsequent steps, including the wash and dissociation procedures, reverse-transcription, and PCR amplification, were carried out as described previously. After reverse-transcription and PCR amplification, the products were purified using the FastGene PCR extraction kit (NIPPON Genetics, Japan). For further selection, the purified DNA fragments were used as insert templates for the HiFi assembly. The vector template for the HiFi assembly was the PCR-amplified product using RD backbone plasmid as the template and the Gibvec-For and Gibvec-Rev primer set. The assembled DNA were then amplified using F1 and Fag-Rev primers to generate DNA templates for additional rounds of ribosome display selection.

In the following rounds of ribosome display selection, we adjusted the binding duration to 1 hour in round 2, 30 minutes in round 3, and 15 minutes in rounds 4 and 5. We also reduced the volume of antibody-binding beads suspension to half in round 4 and to a quarter in round 5.

To monitor the selection efficacy of peptides for the target antibody, Enzyme-Linked Immunosorbent Assay (ELISA) was performed as follows. The DNA template used for the ribosome display described as above was re-amplified with NoSecM-Fw and NoSecM-Rv to remove the SecM sequence. Using the SecM-removed products, the ribosome display-selected peptide pools were synthesized by PURE*frex* 2.1 (GeneFrontier, Japan). 96-well plate (Thermo Scientific, USA) coated with 1% BSA or 1 µg/mL of anti-HA tag antibody in carbonate buffer (pH 9.6) was incubate at 4°C overnight. Then the plates were washed twice with phosphate-buffered saline (PBS) and blocked with 5% skim milk (NACALAI TESQUE, Inc., Japan) at room temperature for 45 minutes. The ribosome display-selected peptide pools were added to the wells and incubated at room temperature for 2 hours. Following this, horseradish peroxidase (HRP)-conjugated T7-tag antibody, diluted 1:3000 in Can Get Signal Solution 2 (TOYOBO, Japan), was added to the plate and incubated at room temperature for 1 hour. After washing the plate three times with PBS-Tween 20, the 1-step Ultra TMB-ELISA Substrate Solution (Thermo Scientific, USA) was added, and the plate was incubated at room temperature for 10 minutes. The reaction was terminated by the addition of 2 mol/L sulfuric acid, and the optical density (OD) was measured at 450 nm using a plate reader (Infinite M200 Pro, TECAN).

### Preparation and Assessment of Next-Generation Sequencing (NGS) Samples

Initially, DNA pools resulting from each round of ribosome display were amplified via PCR using the F1 primer and the uniquely assigned barcode primers Round0/1/2/3-Rv. Subsequently, the PCR products were purified using the FastGene PCR extraction kit (NIPPON Genetics, Japan). Equal quantities of the purified products were then combined in preparation for NGS.

Prior to the sequencing, the quality of the prepared sample was assessed using the High Sensitivity D5000 ScreenTape System (Agilent). Once the sample was determined to be of satisfactory quality, it was pooled together for pair-end Next-Generation Sequencing. This sequencing process was carried out using the Illumina NextSeq550 platform at the Center for Gene Research of Nagoya University.

### Analysis of Next-Generation Sequencing Data and Implementation of Epitope Mapping

The raw data of sequences obtained from Illumina sequencing were initially processed using SeqKit [26] to extract the randomized region of the library. The processed data was loaded onto the MEME motif analysis [27], and consensus amino acid sequence logo was generated.

Then, a series of clean-up procedures were carried out using the R program, which helped in sorting the sequences based on their read counts from each round of ribosome display selection. This step facilitated understanding of the concentration rate of the sequences. Detailed methodologies employed for SeqKit and R program analysis can be found in Supplementary Method 1. Subsequently, the refined amino acid sequence data was loaded into an in-house Python algorithm for epitope mapping analysis (epitope_mapping_2AA.ipynb or epitope_mapping_3AA.ipynb: open at https://github.com/NakanoHideo758/epitope_mapping.git). This process involved examining the frequencies of occurrence for the combinations of two or three amino acids. To evaluate the change in frequency of specific combinations, an enrichment factor (EF) was computed for all possible combinations of two-amino-acid (400 variations) or three-amino-acid (8,000 variations).

The calculation of the EF commenced by determining the occurrence ratio of a particular combination in both pre- and post-ribosome display selections, based on the read counts derived from the NGS data. EF was then calculated using the following formula: EF= (Proportion of the combination post-ribosome display selection)/ (Proportion of the combination in the initial library) Once the EF values for all possible peptide combinations were calculated, each peptide combination was systematically aligned with the potential antigenic protein sequence. For instance, if an antigen sequence is ‘MAAGTQQA---‘, the EF index of the 4th G would be the sum of EFs of the peptide combinations AAG, AGT, and GTQ.

Subsequently, the calculated EF indices were plotted graphically along the amino acid sequence of the antigen. This mapping provided a visual representation of the enrichment of specific amino acid sequences through the ribosome display selection process, thereby identifying potential epitope regions in the antigen that are recognized by the target antibody.

### Alanine Scanning Mutagenesis for the Identification of Binding Epitope of the Anti-HA Tag Antibody

Alanine scanning mutagenesis was performed to determine the binding epitope of the anti-HA tag antibody. Plasmids containing the T7 tag, the HA tag, or site-directed mutated HA tag variants (Y1A, P2A, Y3A, D4A, V5A, P6A, D7A, Y8A, A9S), along with protein D (T20-V109) were constructed and PCR-amplified using F1 and R1 primers, followed by column purification. The purified PCR products were used as templates for cell-free protein synthesis utilizing the PURE*frex* 1.0 kit (GeneFrontier, Japan). The synthesized proteins were then separated using 15% SDS-PAGE and transferred to nitrocellulose membranes (Bio-Rad, USA).

After blocking the membranes with 1% Skim Milk (NACALAI TESQUE, Inc., Japan) at 4℃ overnight, the membranes were washed three times with PBS-Tween 20. The membranes were then probed with either the anti-HA tag antibody (MBL Co Ltd, Japan) to assess binding ability, or the HRP-conjugated anti-T7 tag antibody to measure the quantity of synthesized protein. For the detection of the anti-HA tag antibody, an HRP-conjugated anti-mouse IgG antibody (Fortis Life Science, USA) was utilized as the secondary antibody. The immunoblots were visualized using the 1-step™ TMB-Blotting Substrate Solution (Thermo Fisher Scientific, USA). This process allowed for the identification of specific amino acid residues on the HA tag critical for binding with the anti-HA antibody.

## Results

### Enrichment Analysis of the Model Binary Library

Two model libraries were generated, containing mRNAs of the control-binder (PA tag) and non-binder (Flag tag) at ratios of 1:10 and 1:100. Both libraries underwent a single round of PURE ribosome display selection against the anti-PA tag antibody. The eluted mRNAs from this selection round were then subjected to reverse transcription, converting them back into complementary DNA (cDNA).

The cDNA resulting from one round of selection, along with the cDNA of the PA tag, the cDNA of the Flag tag, and the initial model library (at ratios of 1:10 and 1:100), were all amplified via same PCR. These products were subsequently digested with BamHⅠ and analyzed through DNA-PAGE, as depicted in Figure 2.

**Figure 2.**
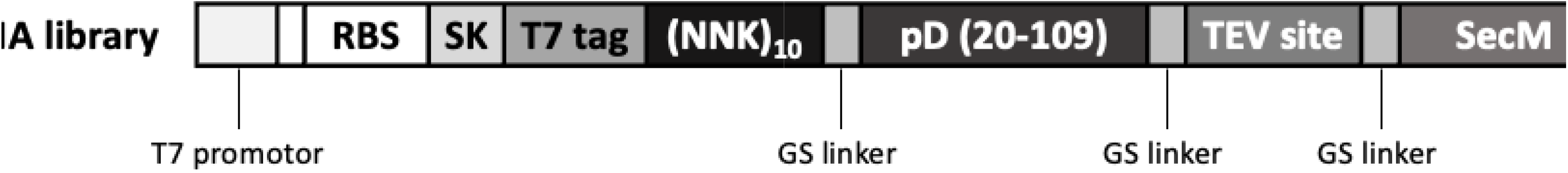
Enriched DNA analysis after one round of ribosome display selection for the model screening library. DNA-PAGE results of BamHⅠ-treated DNA samples from the control binder, non-binder, input DNA library, and the selected DNA pools for both 1:10 and 1:100 binary libraries. C: control binder, DNA samples of PA tag. N: non-binder, DNA samples of Flag tag with BamHⅠ binding site. I: input library, pre-selection DNA samples. S: selected DNA pools, post-selection DNA samples.

The analysis of band intensity demonstrated the successful enrichment of PA tag sequence through the selection process. Band density analysis revealed enrichments of 27 and 17 for the 1:10 and 1:100 model libraries, respectively. These findings strongly indicated a significant enrichment of the PA tag sequence in both libraries following the selection.

### Sequence Analysis of Three Rounds of Ribosome Display Selection: Evaluating Efficiency and Efficacy

Five rounds of PURE ribosome display selection were performed against the anti-HA tag antibody. After each selection round, the DNA pools were transcribed and translated using the PURE system and then analyzed using the Enzyme-Linked Immunosorbent Assay (ELISA). The results, shown in Figure 3, revealed an increase in the ELISA signal during the first three rounds of selection, followed by a decrease in the fourth and fifth rounds.

**Figure 3.**
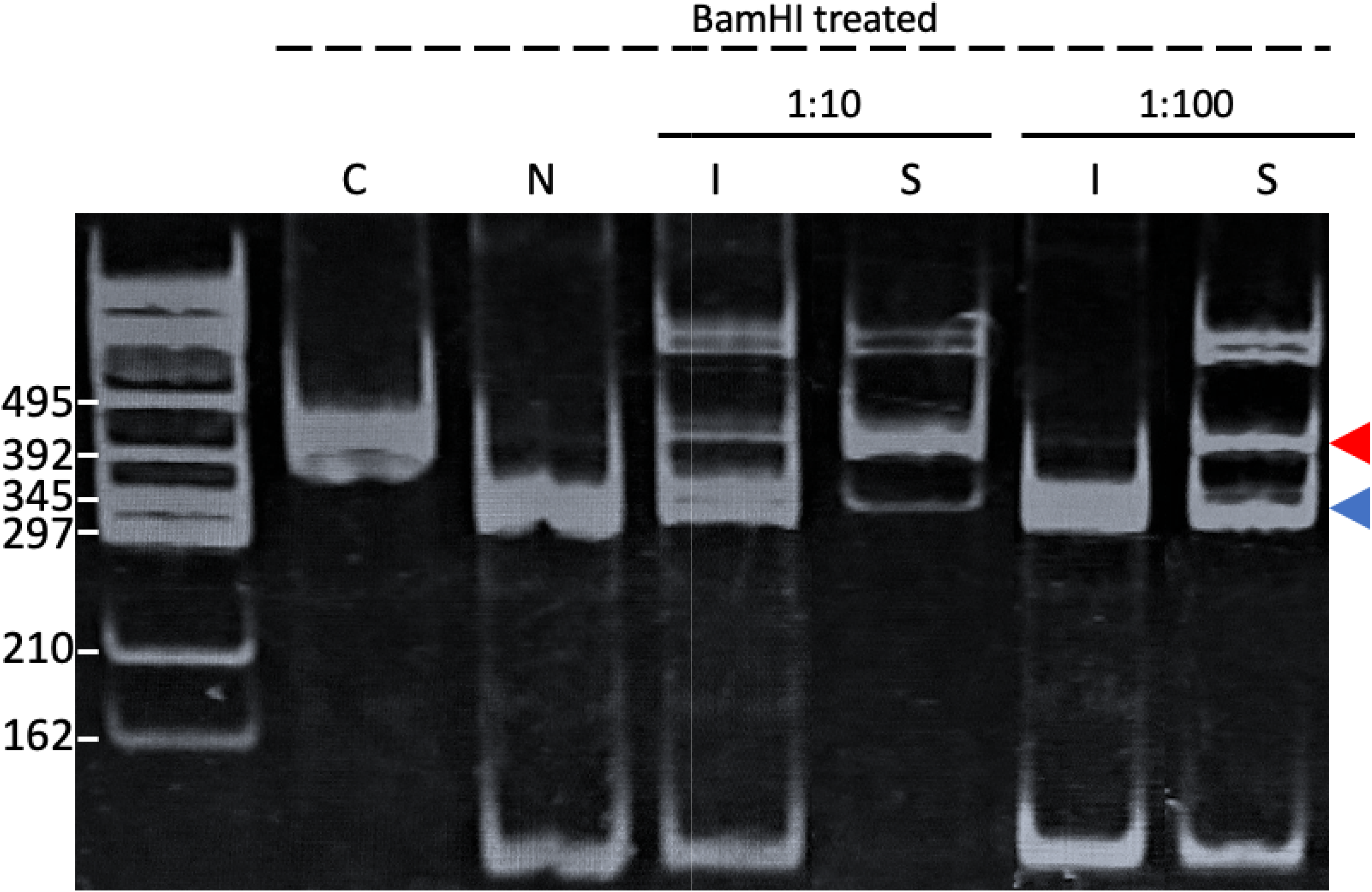
ELISA analysis of pre/post-selection DNA pools. The selected DNA pools and the initial library (R0) were *in vitro* transcribed and translated. ELISA analysis was performed against both BSA and the anti-HA tag antibody.

The DNA pools selected from the initial library and the first three rounds were subjected to Next-Generation Sequencing (NGS) analysis. From the NGS data, the randomized library segment was extracted using SeqKit and further analyzed using the MEME suite for motif analysis. As shown in Figure 4, the consensus sequence from the MEME suite, “LSWMDVPAVS,” included only consecutive three amino acids of the HA tag sequence (YPYDVPDYA).

**Figure 4.**
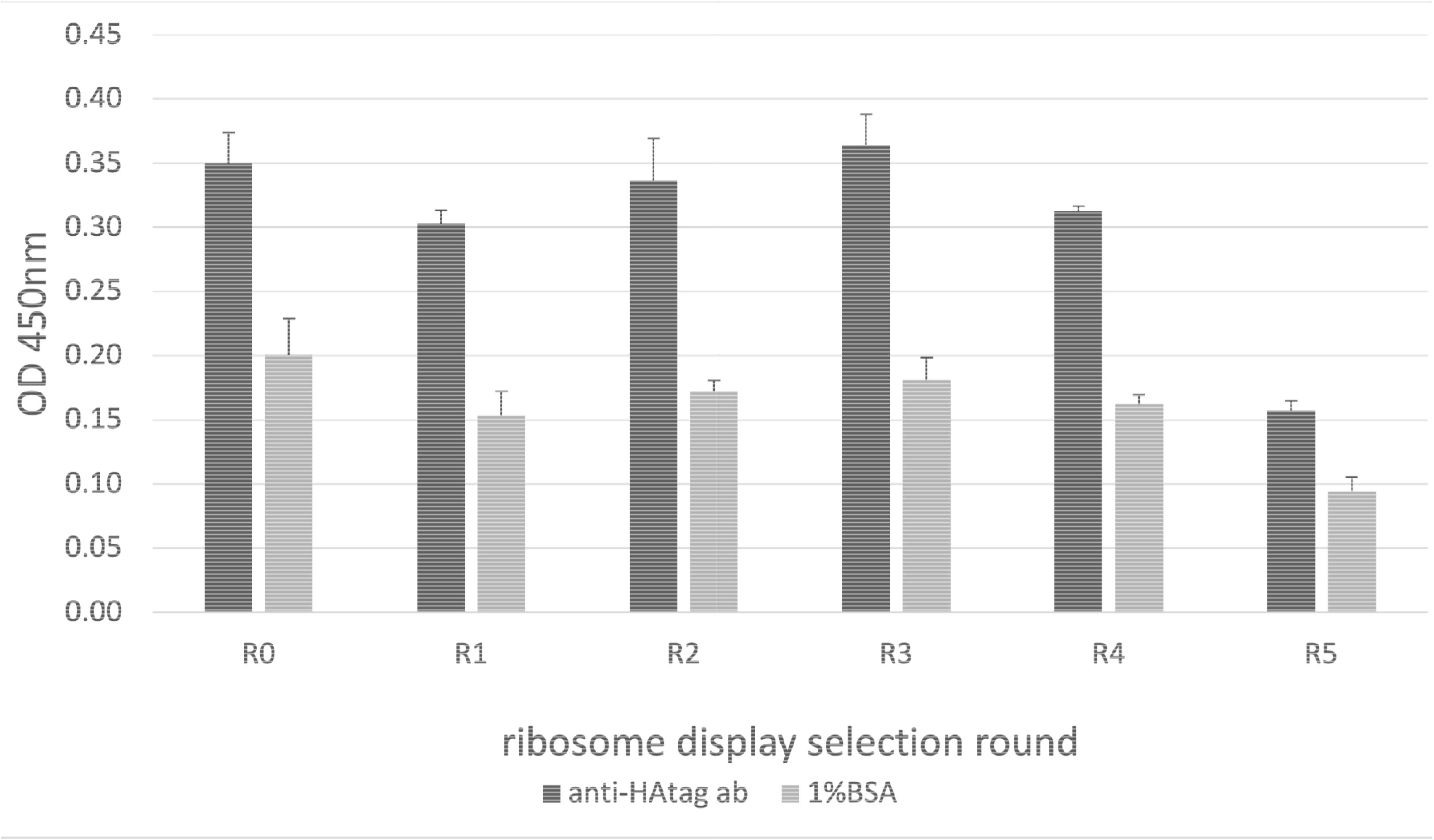
MEME protein motif analysis of the DNA pools from round 1 ribosome display-selected pools.

Further data cleaning and analysis was conducted using the R program, including counting unique sequences and sorting by read counts. The top-ranked sequence, as visualized in Table 1A, matched the consensus sequence identified by the MEME motif analysis. Moreover, the read counts for “LSWMDVPAVS” showed an increasing trend over the three selection rounds.

**Table 1.**
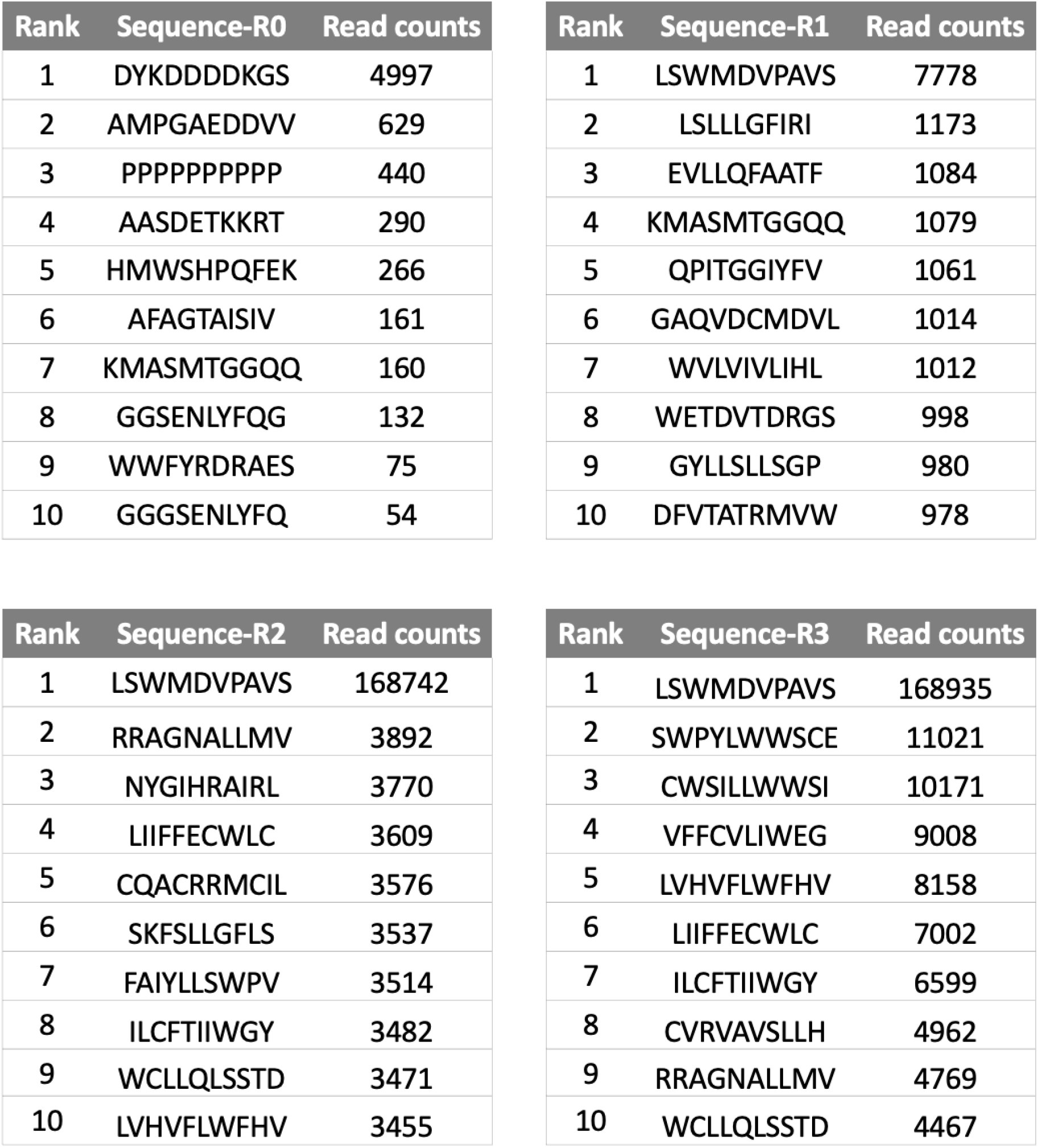
Motif analysis of NGS result. (A) Next-generation sequencing results sorted by read counts of initial library (R0) and selected pools of round 1 to 3 selection. Only the top ten amino acid sequences were displayed. (B) The enrichment rate (ER) and the proportion of “LSWMDVPAVS” following several rounds of ribosome display selection.

The enrichment of the “LSWMDVPAVS” sequence was tracked and presented in Table 1B. While it constituted less than 0.001% of total reads in the initial library, its proportion increased to 0.093% following the first round of selection, indicating a 90-fold increase. This proportion rose even more dramatically in the second round, reaching 12.417%, which was over 130 times higher than the first round. A slight decrease to 11.805% was noted in the third round compared to the second round. Nevertheless, these results showed that the “LSWMDVPAVS” sequence experienced an enrichment of more than 12,000-fold compared to its initial proportion, highlighting the efficacy of the selection rounds in enriching specific sequences.

### Epitope Analysis of Anti-HA Tag Antibody Using NGS Data: Evaluating the Feasibility of Epitope Mapping

To study the binding sites (epitopes) of the anti-HA tag antibody more presicely, we used the obtained NGS data with both R and a specialized Python program we created. We calculated values known as Enrichment Factors (EFs) for combinations of either two or three amino acids. These were then compared to the HA protein sequence, which includes the HA tag that the anti-HA antibody recognizes. An EF value greater than 1 suggests that certain combinations of amino acids are more likely to be the binding sites for this antibody.

Data from the first three rounds were analyzed using a Python program that considered combinations of two amino acids, yielding 400 variations. The analysis from the first round did not indicate significant epitope locations, as the highest Enrichment Factor (EF) was below 2 (see Figure S3). In the second round, a distinct peak was observed for the “DVP” sequence within the HA tag (Figure 5A). However, other observed peaks, such as ‘AVP,’ ‘DV,’ ‘QQ,’ and ‘VP,’ are deemed less trustworthy for epitope identification. This is because genuine epitopes usually involve at least three amino acids. Given that the established HA epitope is “YPYDVPDYA,” the ‘AVP’ sequence is unlikely to be the actual binding site To improve the accuracy of the epitope prediction program, EFs were calculated for all possible combinations of three amino acids, totaling 8,000 variations. These were then aligned with the same antigen sequence. The EF index plot using data enriched in Round 1 (Figure 5B) prominently featured the “DVPDY” sequence with the highest peak. Residues like ‘AVP,’ ‘QQ,’ ‘DV,’ and ‘VP,’ which had high EF index values in Figure 5A, showed lower values in Figure 5B. While other sub-peaks were present in Figure 5B, their EF indices were less than half of the highest peak, “DVPDY.” This suggests that analyzing three-amino-acid combinations provides more reliable results compared to two-amino-acid combinations. Interestingly, the three-amino-acid analysis using Round 1 data (Figure 5B) yielded a distinct peak, contrasting with the two-amino-acid analysis using Round 2 data (Figure 5A). This indicates that even a single round of selection can produce reliable epitope predictions.

**Figure 5.**
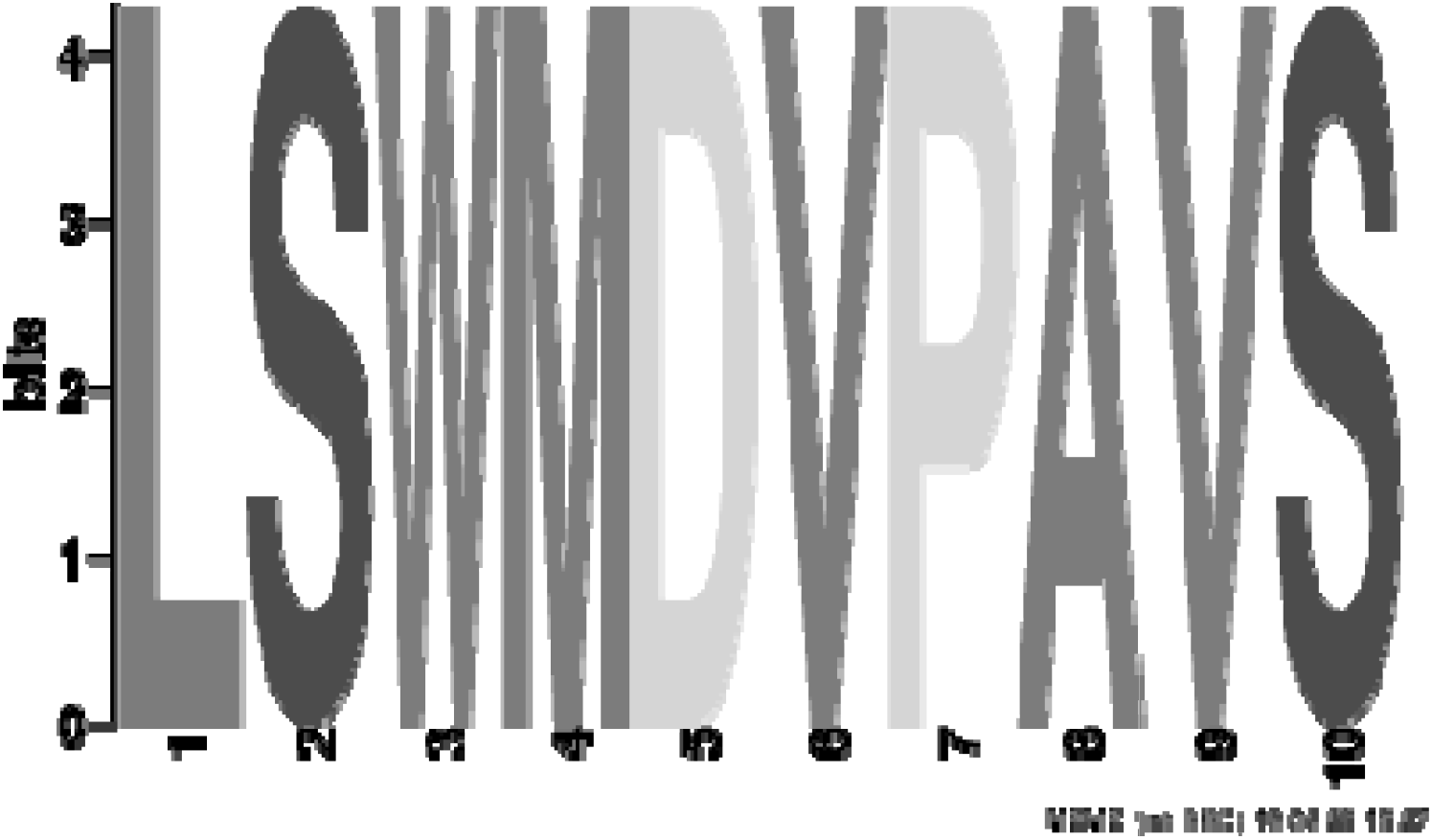
Enrichment Factor (EF) index plot. (A) EF index calculated for combinations of two-amino-acid against hemagglutinin (HA). Residues with high EF values but not part of the epitope were enlarged within a gray frame. The epitope region (YPYDVPDYA) was enlarged within a black frame and highlighted. (B) EF index calculated for combinations of three-amino-acid against hemagglutinin (HA). The epitope region (YPYDVPDYA) was enlarged within a black frame and highlighted. Horizontal axis indicated each residue of hemagglutinin (HA, PDB:3VUN_A). Vertical axis indicated the EF index of each residue.

### Confirmation of the Predicted Epitope for the Anti-HA Tag Antibody

To validate the results of the epitope mapping for the anti-HA tag antibody described above, alanine scanning mutagenesis was performed on the HA tag sequence fused to a protein. The synthesized proteins were then analyzed for their binding to the anti-HA tag antibody using Western blotting.

Figure 6A presented the Western blots of the HA tag-fused protein variants using an anti-T7 tag antibody, providing a quantification of each synthesized protein. Along with the Western blots of the same variants but probed with the anti-HA tag antibody, representing the binding ability of each variant.

**Figure 6.**
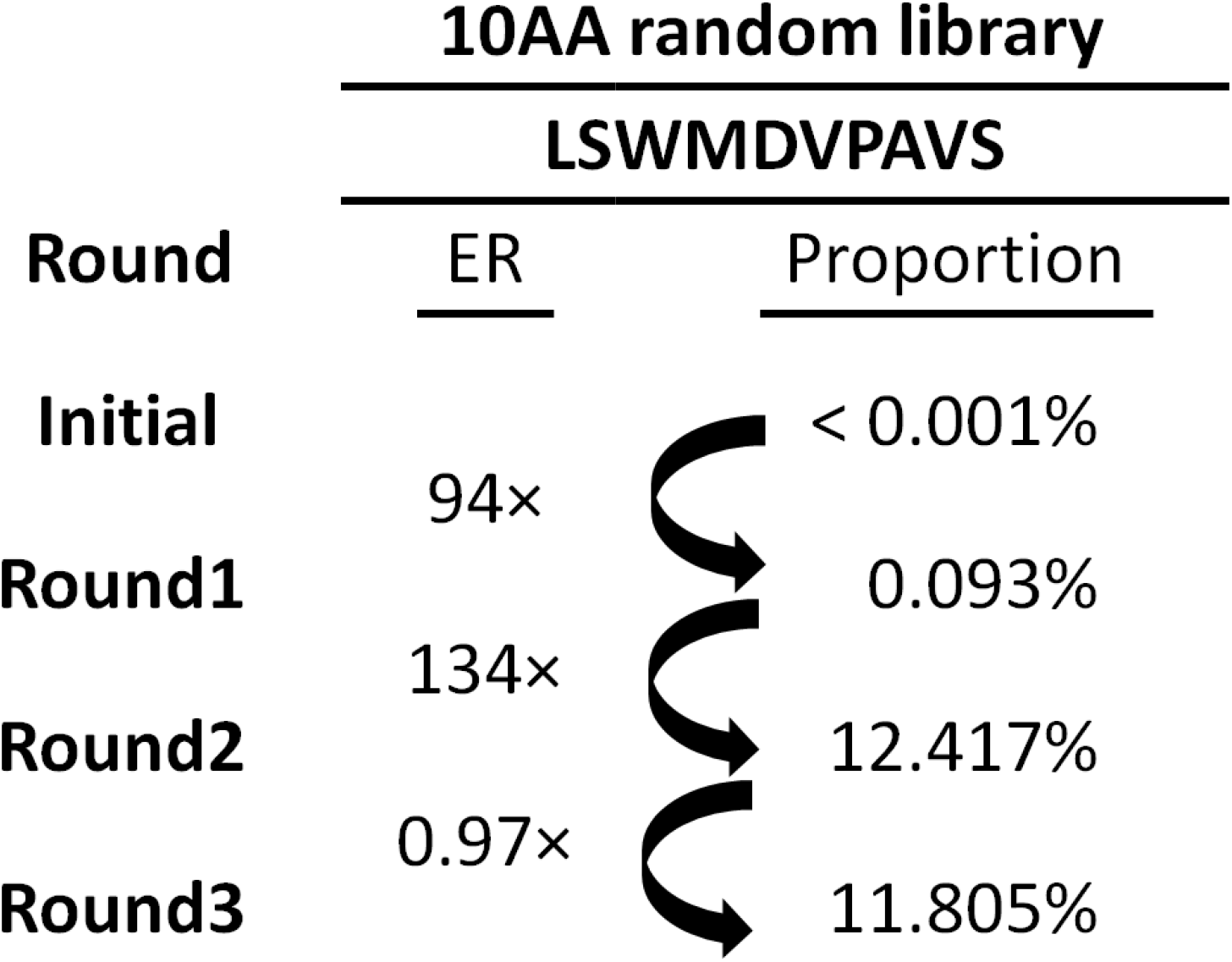
Identification of epitope residues for an anti-HA antibody by alanine scanning. (A) The western blotting results of synthesized quantity and binding ability for all HA variants. For quantity analysis, HRP-conjugated anti-T7 tag antibody was used for detection. For binding ability analysis, the anti-HA tag antibody was employed as the primary antibody for binding, followed by detection using an HRP-conjugated anti-Mouse IgG antibody as the secondary antibody. (B) The relative binding ability of HA variants. Upon mutating residues D4, V5, P6, D7, Y8 to alanine, the binding of anti-HA tag antibody decreased to a maximum of 20% compared to the HA tag. Mutations at other positions did not significantly affect binding. Bars represented the average of triplicate measurements ± SD. Three dots in blue, yellow, and green represented the relative binding ability of each triplicate measurement.

To corroborate the findings in Figure 6A, the binding capacity of each variant was evaluated. Band density for each variant was measured using ImageJ software. The ratio of protein binding to the anti-HA tag antibody was determined by dividing the synthesized protein quantity by the bound quantity for each HA variant. Baseline normalization was carried out using the HA tag as a reference. The relative binding capacity was calculated by comparing the binding ratio of each variant to that of the HA tag. This experiment was conducted in triplicate, and the standard deviation (SD) was computed. The results are presented in Figure 6B.

Among the HA tag (YPYDVPDYA) fused variant proteins, the Y1A and Y3A variants exhibited similar or even higher binding abilities to the HA tag when probed with the anti-HA tag antibody, which means that the change in these positions have almost no effect for the binding with anti-HA tag antibody. For the P2A and A9S variants, a slightly decrease was discovered for the relative binding ability in Figure 6B. However, no detectable bands were found for the D4A, V5A, P6A, D7A, and Y8A variants, indicating these residues were crucial for binding to the anti-HA tag antibody. These findings confirmed that the “DVPDY” residues within the HA tag (YPYDVPDYA) were crucial for binding to the anti-HA tag antibody used in this experiment. Remarkably, this result aligned well with the predicted epitope obtained from the in-house Python algorithm program (Figure 5B), indicating the reliability of this method for epitope prediction.

## Discussion

In this study, we developed a rapid and cost-efficient platform for epitope mapping, utilizing a reconstituted cell-free protein synthesis system, the PURE system, in combination with next-generation sequencing (NGS) and bioinformatics (Figure 1A). The robustness and functionality of this novel PURE ribosome display methodology were empirically demonstrated through a model screening of an anti-PA tag antibody and the subsequent epitope analysis against an anti-HA tag antibody.

Previously, the PURE system has been effectively employed for epitope mapping [16]. This was achieved by creating a stable bond between mRNA, the ribosome, and the protein, facilitated by the SecM arrest sequence [22–23]. In our study, we built upon the previous method developed by Osada and colleagues [16], making few adjustments. We incorporated part of the SKIK sequence, known to promote high-level protein production, along with a T7 tag to aid in detection during ribosome display selection. Recognizing that the SKIK sequence counteracted ribosome stalling downstream [29], we utilized only the SK portion to ensure effective ribosome display in our study. Our modified method proved efficient and feasible in a model screening against the anti-PA tag antibody. Importantly, our method can be executed in a single round within a maximum of 7 hours (including 2 hours for *in vitro* transcription, 40 minutes for *in vitro* translation, 2 hours for ribosome display selection, and 2 hours for the reverse transcription and PCR amplification, as indicated by the red marks in Figure 1A). This considerably shorter timeframe rendered our method significantly faster compared to traditional techniques such as X-ray crystallography.

A key advancement of this study over previous reports of epitope mapping with PURE ribosome display [16, 24] was the integration of NGS and subsequent bioinformatics for the analysis of the ribosome display-selected DNA pools. This contrasted with traditional methods that necessitate individual gene cloning and sequencing. NGS offered distinct advantages, including the high-capacity monitoring of the concentrated DNA sequences throughout a round of selection.

The binding affinity of the ribosome display-selected pools was evaluated using ELISA against the anti-HA tag antibody. As depicted in Figure 3, the initial library exhibited a relatively strong signal against both the anti-HA tag antibody and the negative control BSA. Due to the high background signal in the initial library, we suspected that the present of the T7 tag at the front of the library portion resulted in the non-specific binding to BSA of short peptides with stop codon. These peptides were then detected by the HRP-conjugated T7 tag antibody, leading to the high background signal. Although the background signal decreased after selection, it remained relatively high. This could be attributed to the presence of sequences with stop codons, as indicated by the NGS sequence results, albeit at lower percentages compared to the initial library. The ELISA signal of the anti-HA tag antibody exhibited a gradual increase from the first to the third round of selection, suggesting an enhanced binding capability developed during the ribosome display selection process. Nevertheless, it is worth noting that the difference between the signal of initial library (R0) and the selected results was minimal. This could be attributed to although certain sequences binding to the anti-HA tag antibody during ribosome display process, but they might be masked within a population containing numerous unbound sequences.

The upward trajectory of ELISA signal against the anti-HA tag antibody reached it peal at the third round of selection and subsequently declined during the fourth and fifth rounds. This decrease could potentially be attributed to the emergence of PCR amplification by-products in the fourth and fifth rounds of selection (Figure S2).

The motif analysis of enriched DNA pools, both utilizing the MEME suite and the sorting of read counts utilizing R program, revealed a prominent sequence “LSWMDVPAVS”. Notably, this sequence overlapped with the central portion of the HA tag “YPYDVPDYA” (Figure 4A and Table 1A). The prevalence of this sequence experimented a remarkable 94-fold increase following the first round of selection, and this enhancement escalated to an extortionately 12,000-fold after the second round (Table 1B). However, the third round of selection did not give any further increase, instead showing a slight decrease. This pattern indicated that one or two rounds of selection emerged as more suitable for epitope analysis using PURE ribosome display, as more rounds of selection tended to deteriorate the results of the enriched sequence.

The conventional consensus sequence analysis utilizing the MEME suite or the sorting of read counts suggested that only the three residues of DVP constitute an epitope. However, this appeared unlikely as the HA-tag antibody was expected to have a continuous epitope longer than three amino acids to maintain sufficient specificity as an anti-tag antibody. Hence, we proposed a novel algorithm to predict epitopes based on two main concepts: 1) Calculation of an enrichment factor (EF) for short amino acid sequence combinations, such as two or three amino acids. 2) Alignment of the EF value with the antigen’s amino acid sequence and summation of the overlapping EF values to assign an EF index to each residue in the antigen’s amino acid sequence.

We initially computed the EFs for all two-amino-acid combinations utilizing the NGS data from the initial and second round selected pools (Figure 5A). Although the “DVP” sequence exhibited the highest EF index, other peaks like “AVP”, “DV”, “QQ” were observed, which lied outside the HA tag region.

Subsequently, we calculated the EFs for all three-amino-acid combinations using the NGS data from the initial and first round selected pools. This data was then aligned with the same antigen sequence. The EF index plot displayed a peak at “DVPDY” within the HA tag region “YPYDVPDYA” (Figure 5B). Notably, residues such as “AVP”, “DV”, “QQ” that had shown high EF index in Figure 5A remained low in this analysis. Additionally, the sub-peaks in Figure 5B demonstrated only half the EF index of the highest peak. This substantial difference strongly supports ‘DVPDY’ as the primary candidate for epitope mapping. However, when further analyzing three-amino-acid combinations using the data from the second and third rounds of selection, suboptimal results were obtained. This was because no other peaks were observed except for ‘DVP’ (data not shown). This finding also suggests that a single round of selection is sufficient for reliable epitope prediction, as conducting more rounds of selection may risk the loss of sequence information.

To validate the accuracy of the epitope prediction mentioned above, the epitope of the utilized anti-HA tag antibody was experimentally determined via alanine scanning substitution analysis (Figure 6). All alanine substitutions for the residues “DVPDY” resulted in a decreased band intensity with the anti-HA antibody, while other substitutions did not change substantially. This outcome was in perfect concordance with the EF index analysis obtained via the epitope prediction method using the three-amino-acid combination analysis (Figure 5B).

In conclusion, the application of the PURE ribosome display system enabled the rapid and efficient selection of peptides that bind to a specific antibody within a concise time frame, with a single selection round requiring maximum 7 hours. The integration of NGS analysis facilitated the swift and precise collection of an extensive amount of sequence data, eliminating the need for labor-intensive cloning steps. Moreover, the execution of an epitope analysis program using Python facilitated accurate prediction of consecutive epitope mapping. To further enhance the capabilities of this technology for mapping discontinuous epitopes, additional informatic analysis should be integrated. This could involve utilizing structure prediction tools like AlphaFold 2 [30–31], leveraging the three-dimensional structure of the antigen as a foundation for the analysis.

## Acknowledgement

Jia Beixi gratefully acknowledges the Mizutani Scholarship from the Graduate School of Bioagricultural Science, Nagoya University. The authors express their gratitude to Gene Frontier, Japan, for providing PUREfrex components. Meanwhile, the authors would like to thank Mika Nomoto, Yasuomi Tada, Akiko Akama, and Mikako Yamaguchi for their technical support with next-generation sequencing. This research was financially supported in part by Grant-in-Aid (19H02523 and 22K18919).

---

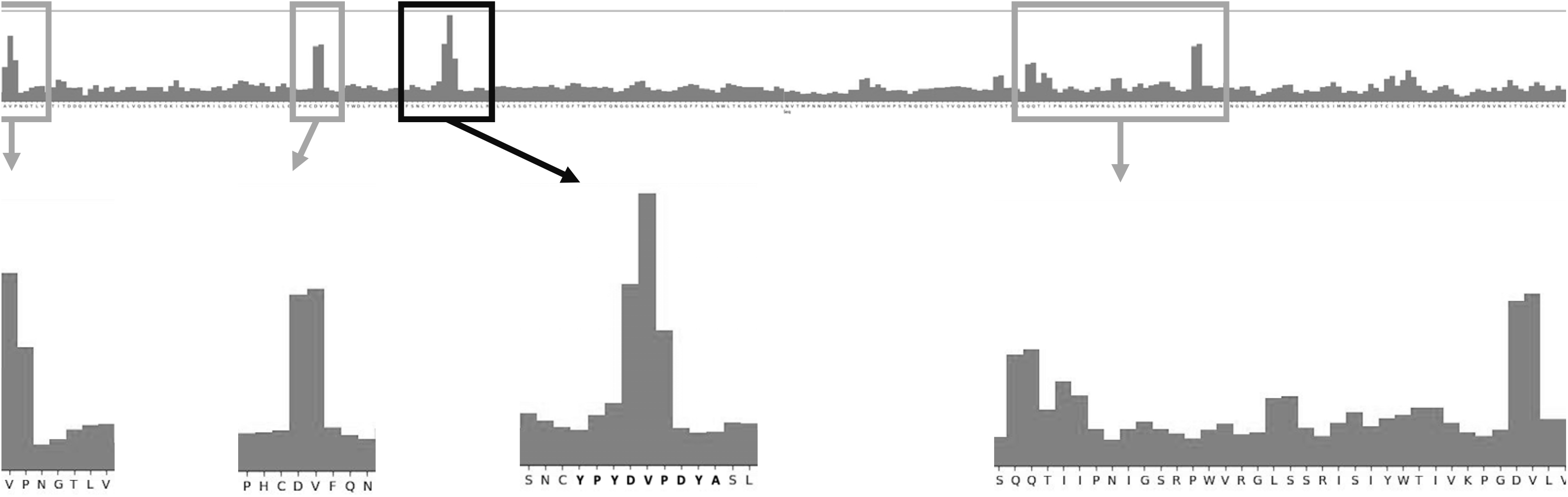

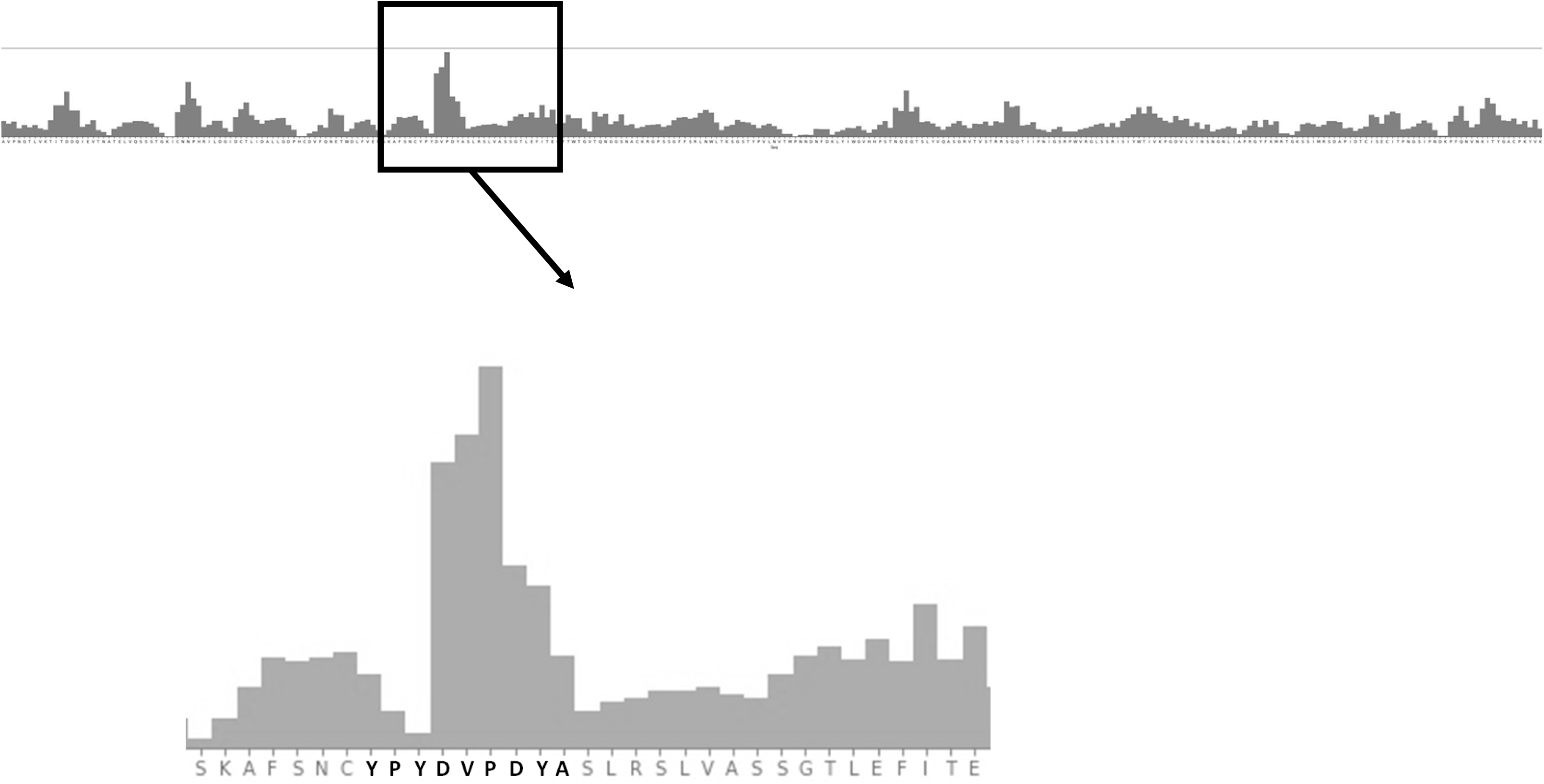

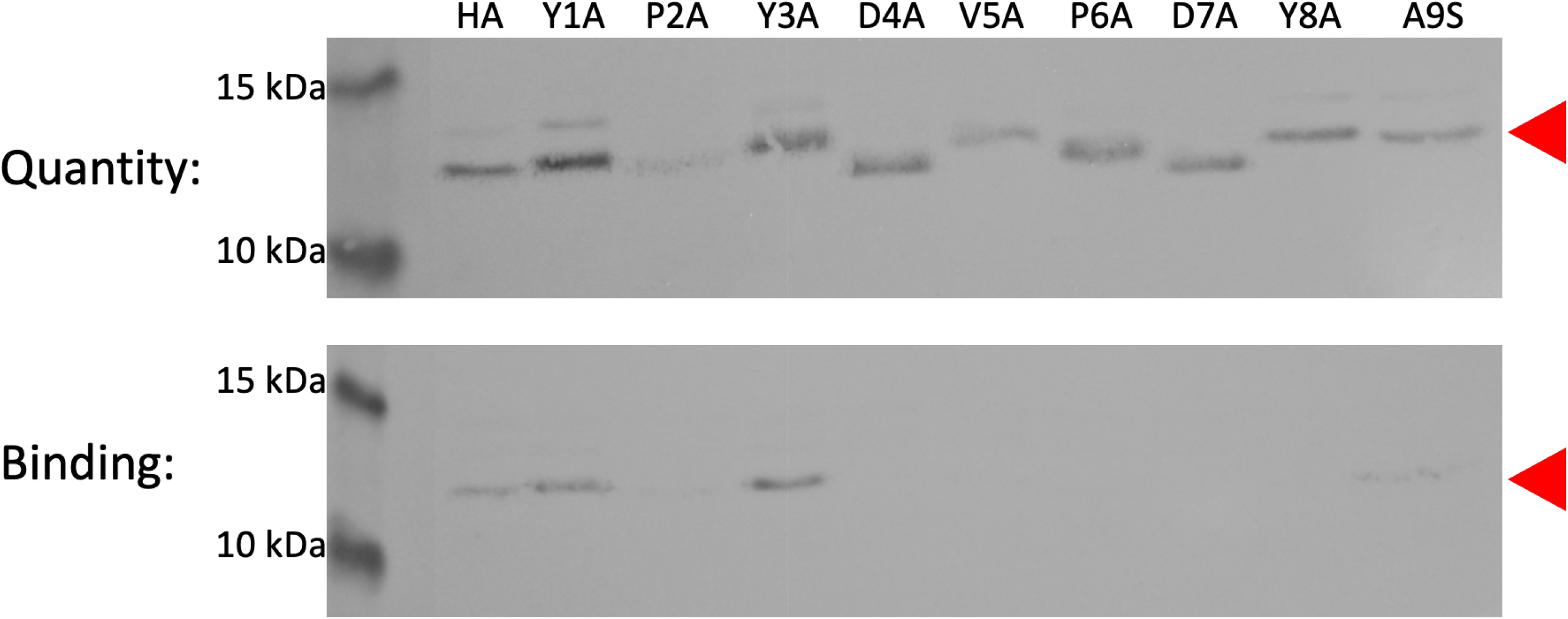

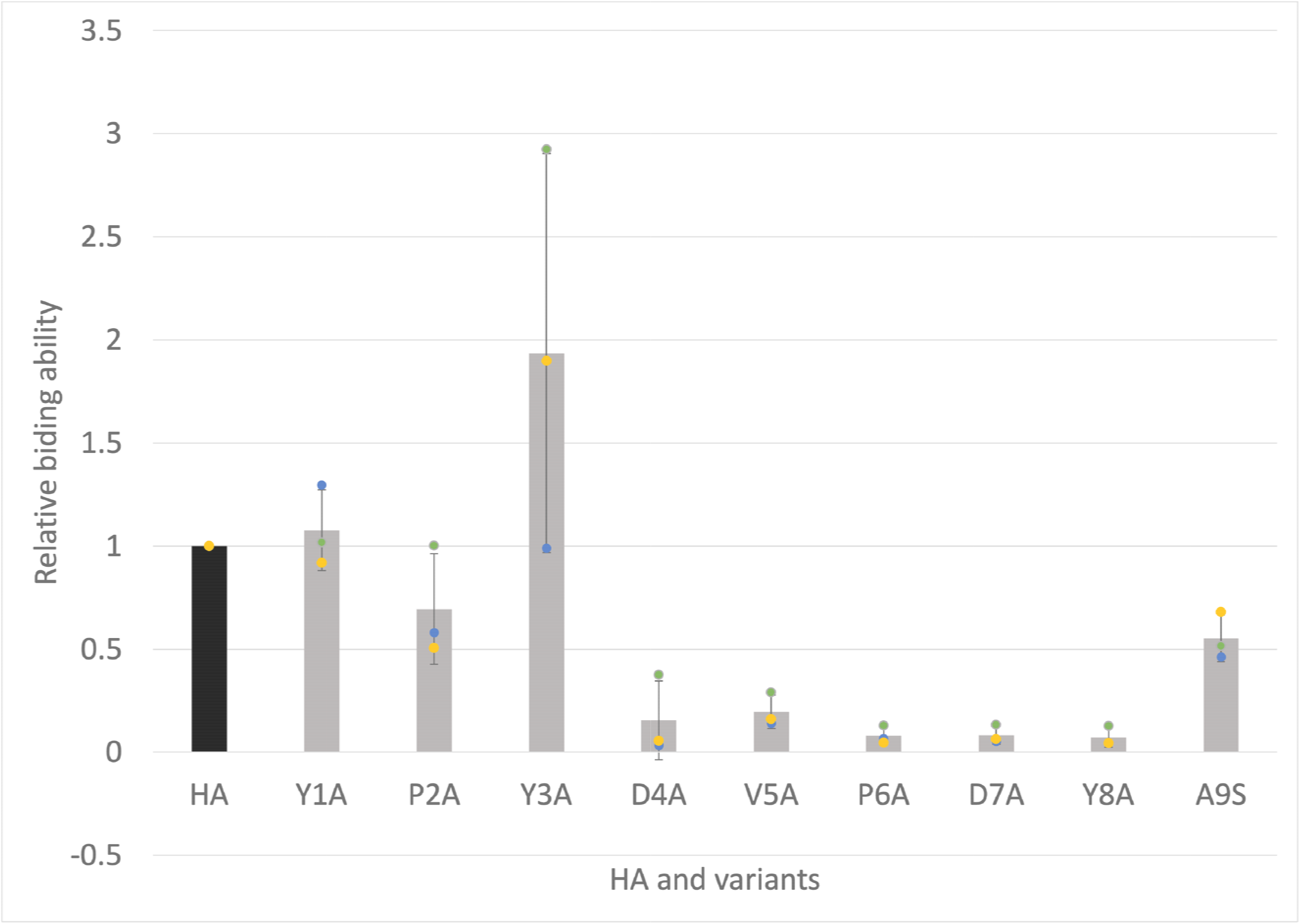

