## Supplementary information for "Rapid and cost-effective epitope mapping using PURE ribosome display coupled with next-generation sequencing and bioinformatics"

**Table S1.** Oligonucleotide sequences. Names and primer sequences during DNA library construction, reverse transcription, and PCR amplification. N=A, T, C, G. K= G, T.

| Name | Sequence (5’→3’) |
|---|---|
| F1 | ATCTCGATCCCGCGAAATTAATACG |
| Fag-Rev | CAGCCGGATCAAGCTTCGAATTC |
| RT-Rv | CATAGTATCGATAGACCTATGCATGCTGGGGTAGAGAATTTC GCAGAACCAC |
| PCR1-Fw | AAGAAGGAGATATACATATGTCTAAGATGGC |
| PCR1-Rv | CATAGTATCGATAGACCTATGCATGCT |
| PCR2-Fw | AAGAAGGAGATATACATATGTCTAAGATGGCTTCGATGACTG |
| PCR2-Rv | GGGGTAGAGAATTTCGCAGAACCACCACCACCCT |
| Libvec-For | GTGGTGGTTCTACCGCGACCGC |
| Libvec-Rev | AGAGCCTCCCATCTGTTGGCCACC |
| 10AA-Fw | GTGGCCAACAGATGGGAGGCTCTNNKNNKNNKNNKNNK NNKNNKNNKNNKNNKGGTGGTGGTTCTACCGCGACC |
| Lib-Rev | GGTCGCGGTAGAACCACC |
| Gibvec-For | TCTGCGAAATTCTCTACCCCGGTTTG |
| Gibvec-Rev | ATATGTATATCTCCTTCTTAAAGTTAAACAAAATTATTTCTAG |
| R1 | TCCGGATATAGTTCCTCCTTTCAG |
| Round0-Rv | GGGCTTTCGGCGCGGTCGCGGTA |
| Round1-Rv | TTTAGGGCGGCGCGGTCGCGGTA |
| Round2-Rv | CCCGAAACGGCGCGGTCGCGGTA |
| Round3-Rv | AAATGGGCGGCGCGGTCGCGGTA |

**Figure S1.** Schematic representation of the control binder (PA tag) and Non-binder (Flag tag) DNA templates used for model screening of ribosome display.

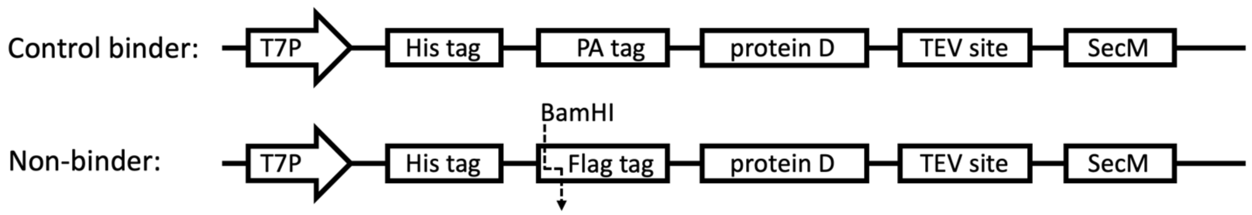

**Supplementary method 1.** The data analysis of NGS results performed using Seqkit and R program.

Seqkit:

```
$ seqkit fq2fa result.fastq > result.fasta
$ seqkit grep -sp barcode sequence result.fasta l seqkit seq -w > RoundX_R.fasta
$ seqkit subseq -r 35:64 RoundX_R.fasta > RoundX_R_seq.fasta
$ seqkit seq -rp RoundX_R_seq.fasta > RoundX_seq.fasta
```

R program:

```
library(seqinr)
RoundX_seq<- "RoundX_seq.fasta"
result_RoundX<- read.fasta(RoundX_seq, seqtype = "DNA")
AA_RoundX<- lapply(result_RoundX, function(x){translate(x, ambiguous = T)})
out_RX<- "RoundX_AA_seq.fasta"
RoundX<- read.table("RoundX_AA_seq.fasta", header = F)
Round0_fre<- read.csv("frequencytable_Round0.csv")
frequencyX<- table(RoundX_lib_seq)
Order_RoundX<- sort(frequencyX, decreasing = T)
write.csv(Order_RoundX, "frequencytable_RoundX.csv")
library(readr)
RoundX_fre_seq<- RoundX_fre[,-1]
Nostar_X<- strsplit(as.character(RoundX_fre_seq$RoundX_lib_seq),"")
judgeX<- numeric()
for (i in 1: nrow(RoundX_fre_seq)) {
  judgeX[i] = length(which(Nostar_X[[i]] %in%c("X","*"))>0)
}
sum(judgeX)
nostar_X<- RoundX_fre_seq[judgeX==0,]
write.csv(nostar_X, "NostarX_RoundX.csv")
```

MEME motif analysis:

```
meme library.fasta -protein -nmotifs 5 -o Round1_AA.out
```

**Figure S2**. Agarose gel electrophoresis after PCR amplification of DNA from initial library and ribosome display-enriched DNA pools. R0 represented the initial library, R1-R5 represented the ribosome display-enriched DNA pools of Round1 to Round5 selection. Red triangle indicated the position of amplified DNA bands. Blue triangle indicated the by-product of PCR amplification.

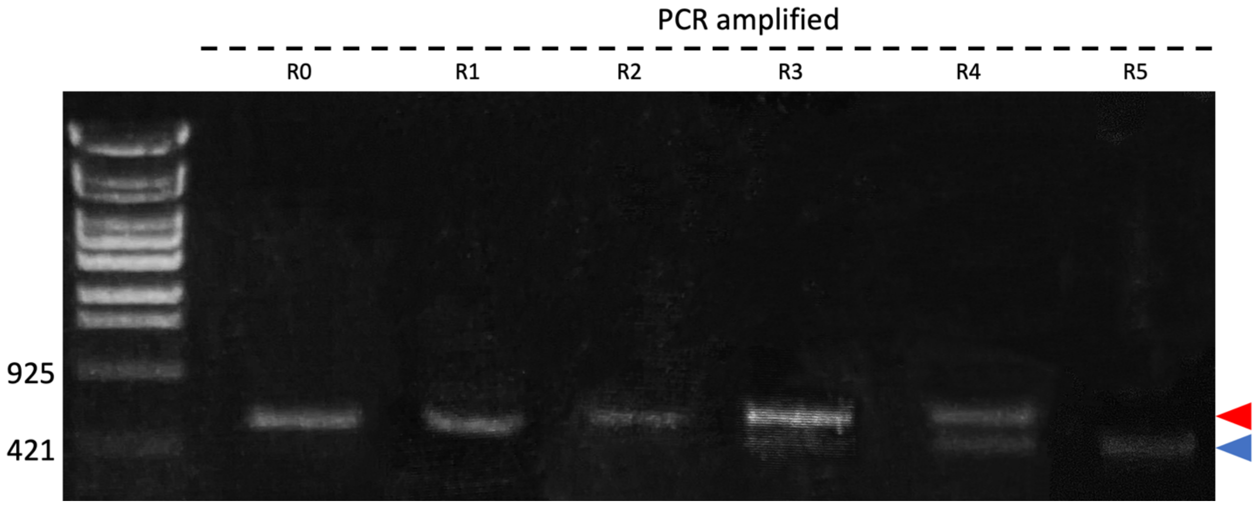
